## Supplementary figures and images for "Male-Biased Cyp17a2 Orchestrates Antiviral Sexual Dimorphism in Fish via STING Stabilization and Viral Protein Degradation"

### Figure S1

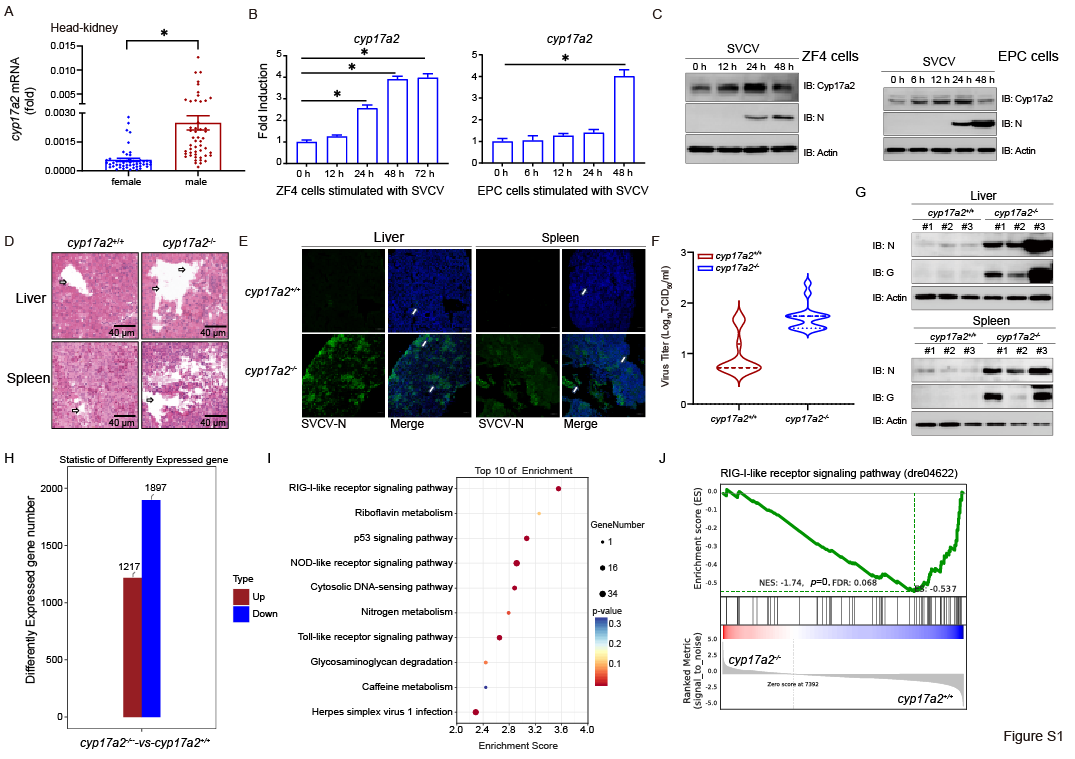

### Figure S2

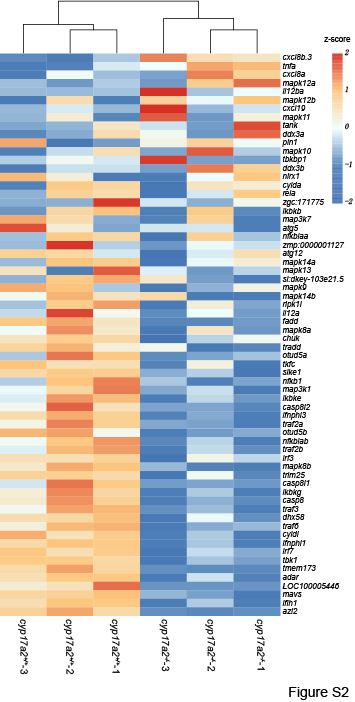

### Figure S3

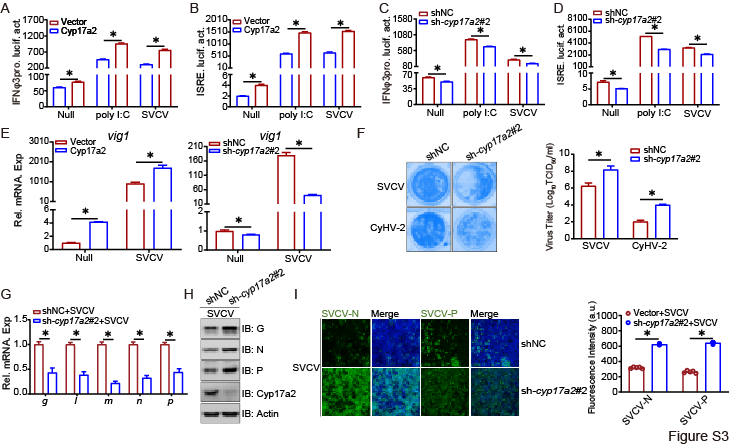

### Figure S4

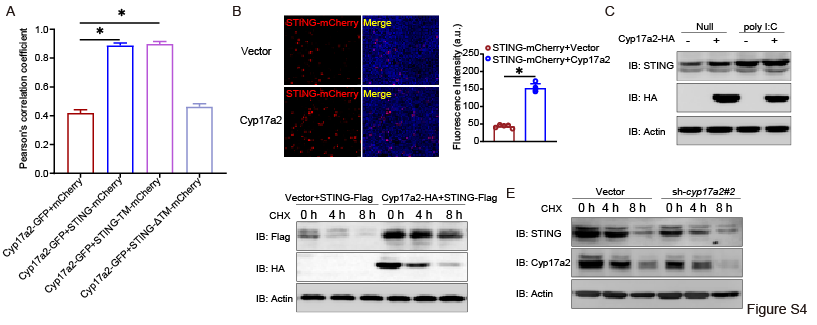

### Figure S5

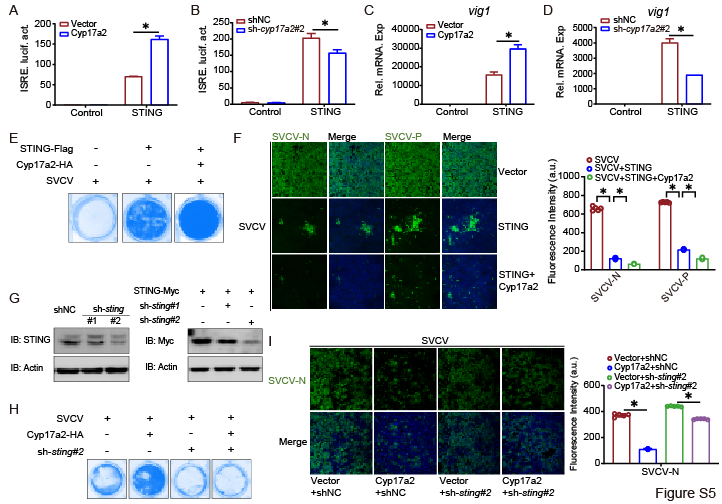

### Figure S6

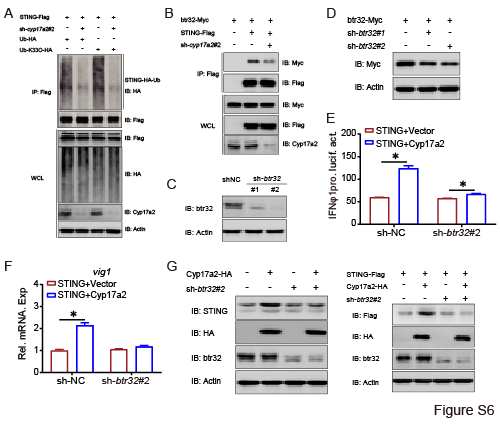

### Figure S7

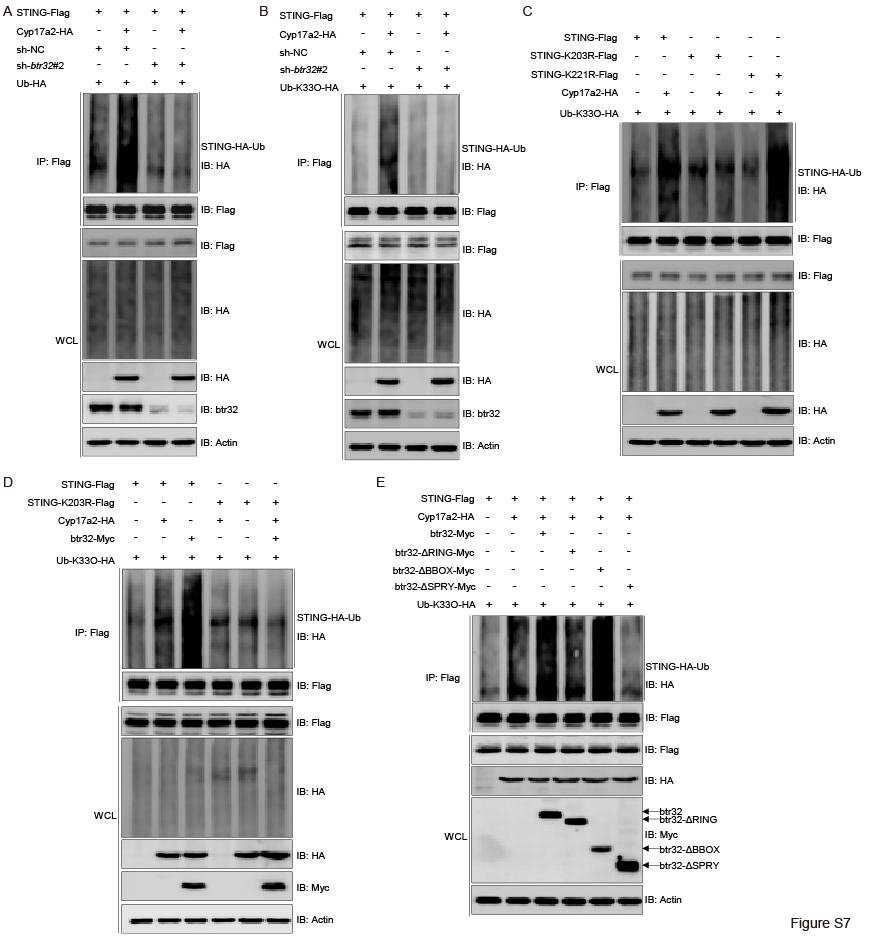

### Figure S8

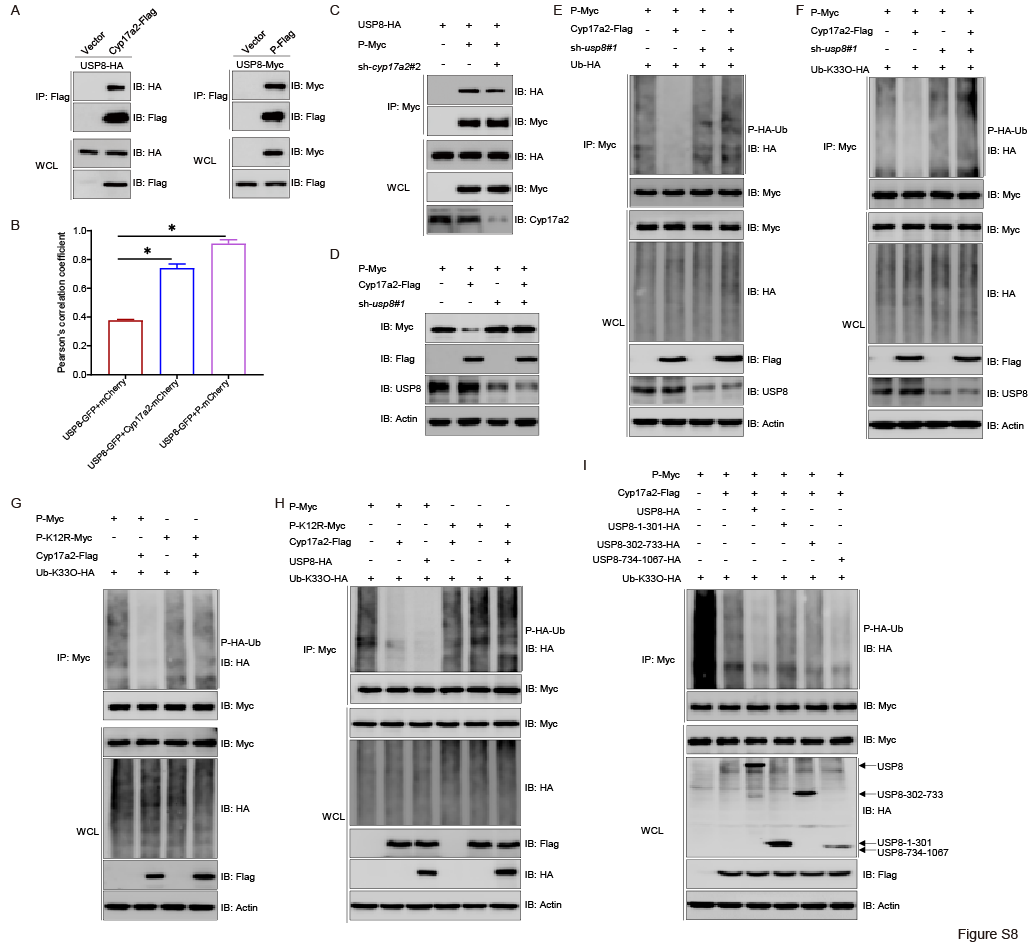
